# Effect of changing the direction of the force vector of the mandible in relation to the skull base: study in rabbits

**DOI:** 10.1101/2022.10.02.510542

**Authors:** Milton Cristian Rodrigues Cougo, Edela Puricelli, Alexandre Silva de Quevedo, Luciana Santa Catarina, Felipe Ernesto Artuzi, Deise Ponzoni

## Abstract

The temporomandibular joint has a great capacity for functional adaptation. The aim of this study was to evaluate, bilaterally, the influence of unilateral modification of the direction of the mandibular force vector in relation to the skull base in rabbits. Thirty New Zealand rabbits (Oryctolagus cuniculus L.) were randomly divided into two groups (n = 15/group): test (modification of the mandibular force vector) and control (no modification of the mandibular force vector). The animals were killed at 20, 40, and 60 days postoperatively. Histomorphometric evaluation of the temporal and condylar joint structures bilaterally always showed significant differences (P < 0.05) between the test and control groups on both sides of the TMJ. The results demonstrate that the rabbit temporal bone and mandibular condyle showed significant adaptive capacity as a biological response to mechanical forces on both the operated and opposite sides.

## INTRODUCTION

The temporomandibular joint (TMJ) is a complex synovial articulation. It is characterized as a bilateral diarthrosis, comprised between the mandibular condyle, the articular fossa, and the articular eminence of the temporal bone [1].

The TMJ differs from other joints in that it is not covered by hyaline cartilage, but by a layer of avascular fibrous connective tissue. According to Okeson (1992) [2], this tissue has two significant factors in TMJ function: greater resistance to wear with aging and greater regenerative capacity. Adaptation and remodeling are the main biological mechanisms responsible for maintaining the morphological and functional balance of the joint. In adulthood, with reduced growth function, remodeling, although with a lower quantitative response, is considered the most important mechanism for maintaining functional balance [3].

According to Puricelli (1997) [4], in the human TMJ, the functional force vector in the mandible has a posterior-anterior/inferior-superior direction through the condyle to the articular tubercle of the zygomatic process. Upon completion of growth, the influence of dynamic forces applied to the tissues is maintained by continuous stimulation of apposition and resorption. Mandibular movements can be continuous or intermittent, can act alone or in combination, and result from the static and dynamic loading of the TMJ [5,6].

Mechanical loading of the TMJ is necessary for the growth, development, and maintenance of joint tissue. Generally, dynamic loading provides an anabolic effect on the joint tissues, while static loading, especially if prolonged or excessive, induces a catabolic effect. As different movements occur simultaneously between the joint surfaces, the TMJ is subjected to a combination of loads, such as compression, tension, and shear [7]. Thus, the layers of articular cartilage and the fibrocartilaginous disk can undergo deformations, which are accompanied by internal tissue changes. Therefore, evidence confirms that the mandibular condyle and glenoid fossa have ample capacity for functional adaptation [6,8,9].

Remodeling of the glenoid fossa and compensatory growth of the mandibular condyle adjust with the anatomical position of the condyle in the fossa. The subarticular proliferative zone can support anabolic and catabolic bone remodeling to change the shape and position of the temporal fossa in response to environmental changes [6,8,9]. The role of the condyle in the growth and development process of the TMJ and its adaptive response to mandibular advancement are known to be notably greater, particularly during the period of accelerated growth, when compared to the glenoid fossa [6,8,10]. However, although the patterns of adaptive responses are different in both structures, they act together in a harmonic manner [11]. Therefore, mandibular growth modification is the result of morphological cell differentiation and hypertrophic changes in the condyle and glenoid fossa [8].

The purpose of this study is to evaluate in rabbits, bilaterally, the influence of unilateral modification of the direction of the mandibular force vector in relation to the skull base.

## MATERIALS AND METHODS

### Experimental Design

Thirty male New Zealand rabbits (*Oryctolagus cuniculus L.*), 4 months old, weighing between 3 and 4 kg were used in this study. The rabbits were housed in individual cages at a temperature of 20 ± 1°C and relative humidity of 40-60%, air exhaust system and light: dark cycle of 12/12 hours. Feed (standard for the species), cabbage, banana, and water were provided *ad libitum*. This study was conducted in accordance with Law no. 11,794/2008, which establishes the standards for the use of animals in research in Brazil and was approved by the Ethics Committee on Animal Use (CEUA) of Hospital de Clínicas de Porto Alegre (HCPA). All rabbits used in the study came from the same supplier, who is registered at HCPA. The health conditions of the rabbits were monitored throughout the study by a veterinarian.

The rabbits were randomly divided into two groups: a test group (GT/n = 15) and a control group (CG/n = 15). Each group was subdivided into three subgroups according to the time of death of the rabbits (20, 40, and 60 days postoperatively). As follows: GT20/n = 5, GT40/n = 5, GT60/n = 5, CC20/n = 5, CC40/n = 5, and CC60/n = 5.

The surgical procedure was performed on the right mandible of all animals. For the test group (GT/n = 15), the surgical intervention caused a change in the direction of the mandibular force vector in relation to the skull base. For the control group (CG/n = 15) the bone segments were fixed in their original position (without changing the direction of the mandibular force vector relative to the skull base).

### Surgical procedure

Rabbits were premedicated with ketamine (15 mg/kg), midazolam (1 mg/kg), and morphine (2 mg/kg) intramuscularly. General anesthesia was supplemented with isoflurane 5% via inhalation. After trichotomy, skin antisepsis was performed using a 0.5% alcoholic chlorhexidine solution. Then, the skin incision was made with a number 15 scalpel blade and number 3 handle, in the rabbit’s right submandibular region. The curved blunt-point scissors allowed tissue divulsion by planes. After incision of the periosteum and detachment, access to the cortical bone in the right mandibular angle was achieved.

After hemostasis and ample visualization of the region, a titanium microplate was positioned distal to the mandibular angle and parallel to the long axis of the mandibular body. With the aid of curved mosquito forceps, the fixation was performed to define its initial position, and with a ½ spherical steel bur four points were defined, corresponding to the four holes of the microplate.

After horizontal perforations corresponding to plate fixation, cortical bone perforations were made in the mandibular angle region, determining the vertical fracture line to the long axis of the mandibular body, followed by cortical ostectomy with complete continuity solution and fracture under constant irrigation with distilled water using a cylindrical drill (13mm × 6mm). The fractured proximal mandibular segment obtained was composed of the posterior portion of the ramus and condyle (Fig. 1A).

**Fig. 1.**
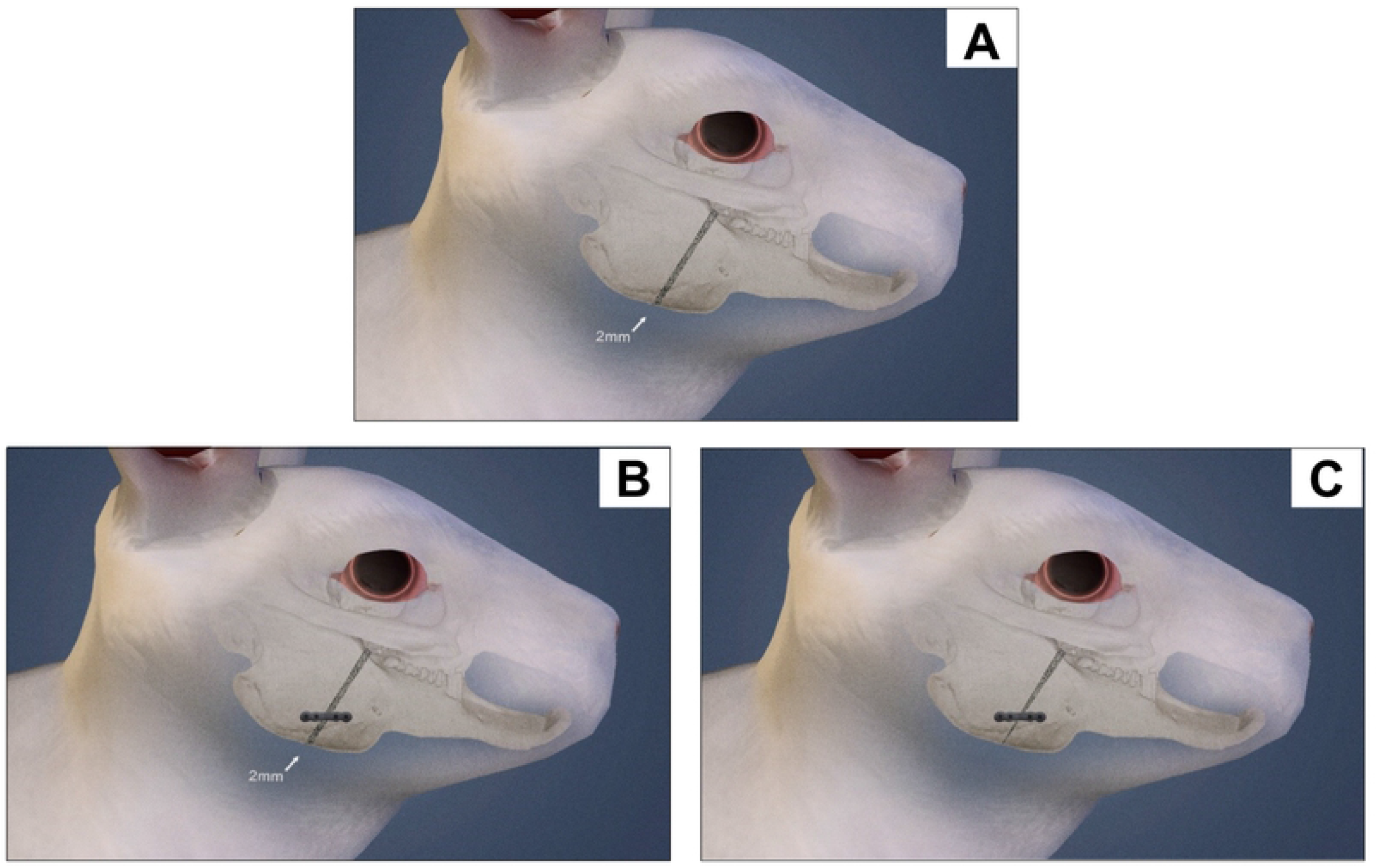
Schematic drawing of the rabbit. **A** - Ostectomy area in the mandible, right side. **B** - Fixation of the bone segments in the original position, with microplate and micro screws, in the control group. The arrow indicates the maintenance of a 2 mm space between bone stumps throughout the vertical ostectomy, without changing the direction of the force vector of the mandible in relation to the skull base. **C** - Fixation of bone segments with microplate and micro screws in the test group. The arrow indicates the approximation of bone stumps in the basilar region, changing the direction of the vector of force of the mandible in relation to the skull base.

Then, it was removed, and from the approximation of the bone stumps in the basilar region, the definitive position of the plate was determined, locating the proximal screws at 2 millimeters in the same direction, in linear continuity of the original position.

The microplate was fixed with micro screws in the distal segment and then in the proximal segment. In the control group, the bone fixation maintained the original position with 2 mm between the bone stumps (Fig. 1B). The new position of the proximal segment provided a sagittal and medial tilt of the articular condyle, displacing it in the anteroposterior direction (Fig 1C). The condyle now articulates in the region of greater temporal convexity, changing the direction of the mandibular force vector on the TMJ.

The osteosynthesis system used was manufactured specifically for the study. It was composed of commercially pure titanium microplates and micro screws, with the following composition: Ti: Bal: N: 0.05 ppm; h: 18.00 ppm; Fe: 0.25 ppm; O2: 0.11 ppm. Each microplate had the following characteristics: four holes with space; distance between holes (center to center) of 2.5 mm; gap distance (center to center) of 4 mm; width of 2.5 mm; and thickness of 0.5 mm. Each micro screw had a thread diameter of 1.5 mm; thread pitch of 0.5 mm; active thread length of 2.4 mm; screw head diameter of 2.4 mm; and a double-slotted, contraposed lateral screw clamping system.

After irrigation of the surgical wound with distilled water, the tissues were sutured with single isolated stitches with 4-0 polyglactin thread in the deep layers and 5-0 mononylon in the dermis.

### Histological preparation

Following the schedule of 20, 40, and 60 postoperative days, the animals were killed according to the following protocol: pre-medication with ketamine (15 mg/kg) and midazolam (1 mg/kg) IM and deep anesthesia with propofol injection, IV, at a dose of 7 mg/kg; followed by injection of 10% potassium chloride, at a proportion of 1 ml/kg of weight. After death and tissue dissection, hemi section of the rabbits’ skulls was performed, and the right and left temporomandibular joints were removed. The pieces were fixed in 10% buffered formalin for two weeks and then decalcified in 10% citric acid for five days. After the decalcification process, they were sectioned in the sagittal plane, dividing them into two fragments, one medial and one lateral. From the center, 0.4μm sagittal sections were made. The slides were prepared in hematoxylin/eosin (HE) stain for histological analysis.

### Quantitative Analysis

Images were captured randomly at 400× magnification by the blinded evaluators using a camera attached to an Olympus Optical Co. binocular microscope (model U-LH100-3) and an AOC computer (model 2236 Vwa), using Qcapture Pro software (version 5.1; Quantitative Imaging Corporation, Inc.; 2005). A polarized light microscope was used.

#### Temporal bone

The thickness was measured in the anterior, middle, and posterior regions of the temporal structure. A vertical line extending from the surface of the temporal bone to the limit of immature bone (delimited by mature bone - Havers System) was used as reference (Fig. 2).

**Fig. 2.**
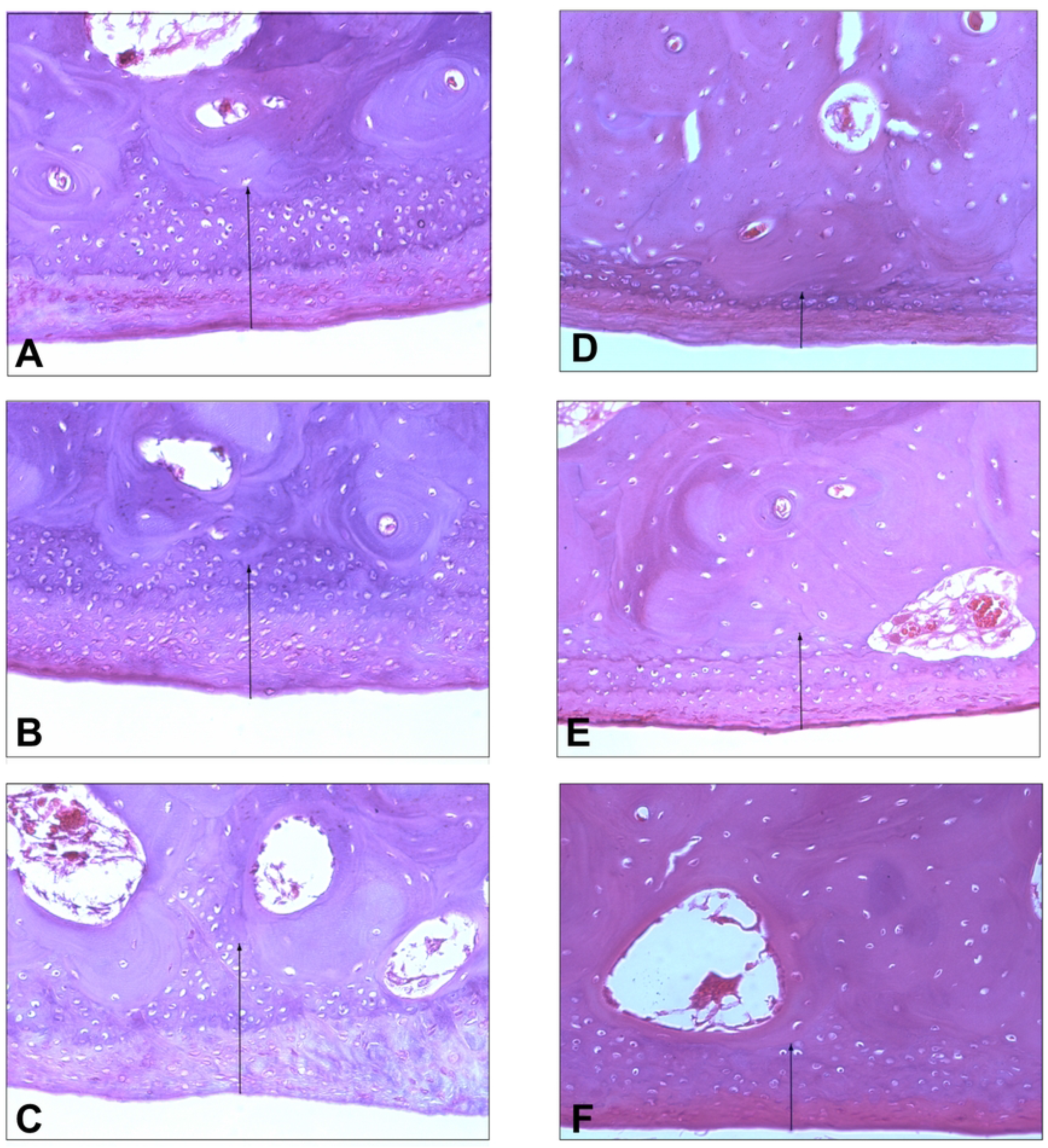
Histological images of the temporal bone at 60 days. Test group, in the anterior (A), middle (B), and posterior (C) regions. Control group, in the anterior (D), middle (E), and posterior (F) regions. The arrows indicate the measurement regions. Staining: hematoxylin and eosin (H & E), original magnification 400X.

#### Mandibular condyle

thickness was measured in the anterior, middle, and posterior regions of the condylar structure. A vertical line extending from the condylar surface to the ossification zone was used as reference.

**Fig. 3.**
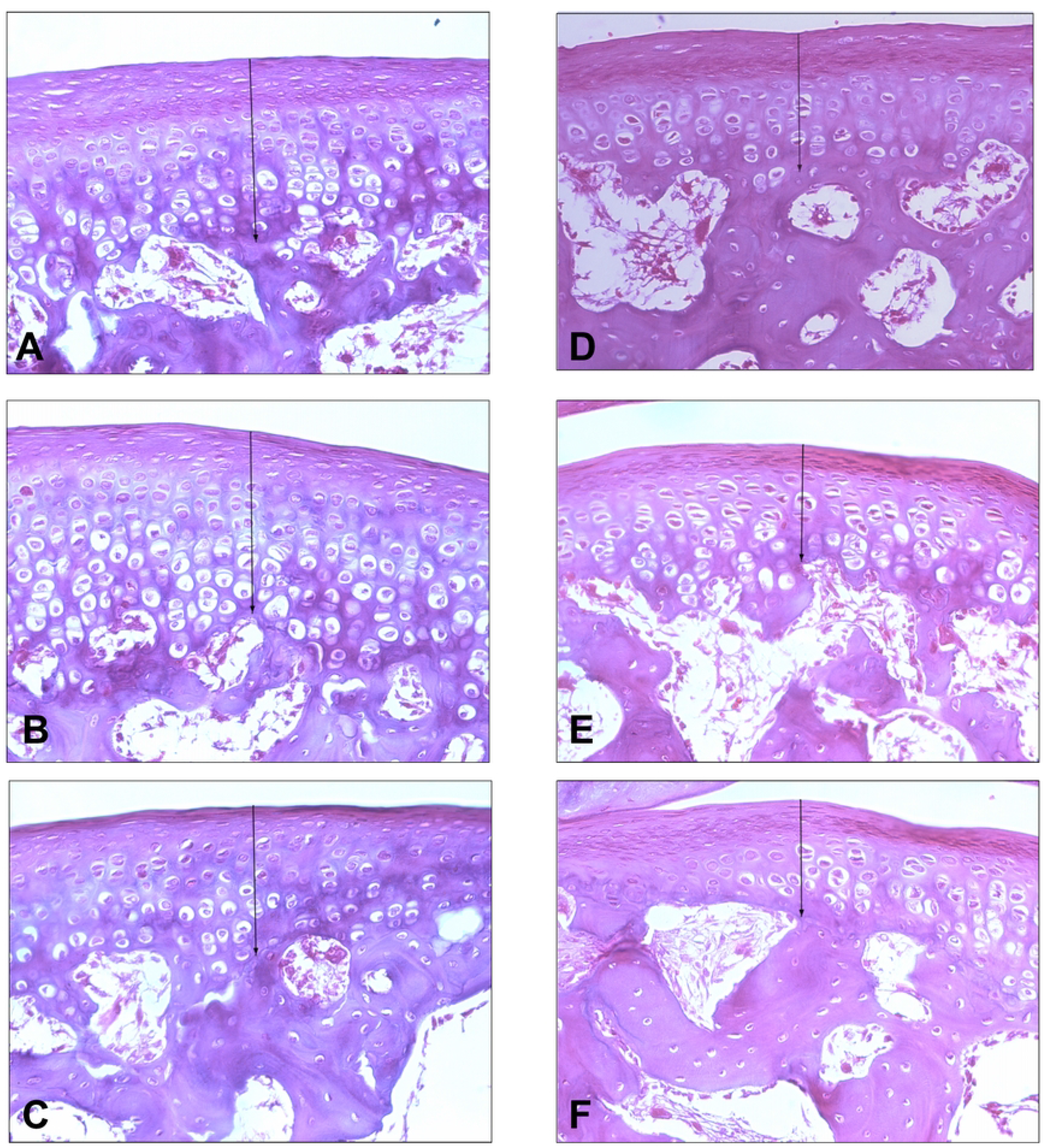
Histological images of the condyle at 60 days. Test group, in the anterior (A), middle (B), and posterior (C) regions. Control group, in the anterior (D), middle (E), and posterior (F) regions. The arrows indicate the measurement regions. Staining: hematoxylin and eosin (H & E), original magnification 400X.

### Calibration

The measurements were performed by two blinded evaluators. The Bland-Altman test showed 95% agreement between the measurements.

In the measurements obtained with the quantitative analysis of the temporal bone, the mean difference between the evaluators was −0.038cm. The limits were between −0.295 and 0.219.

For the measurements obtained with the quantitative analysis of the mandibular condyle, the mean difference between the raters was −0.34 cm. The limits were between −0.187 and 0.118.

### Statistical Analysis

The data were entered into the Excel program and later exported for statistical analysis to the SPSS Version 20.0 program. The measurements of the temporal bone and condyle in the anterior, middle, and posterior regions were presented by mean and standard deviation. The comparison between control and test at 400× magnification; postoperative times (20, 40, 60 days); and sides (right and left TMJ) was performed by the Bonferroni-adjusted Generalized Estimating Equation Model (GEE) method. The total value, without stratification of regions, was described by means and standard deviations and compared by Student’s t test for independent samples. A 5% significance level was considered for the established comparisons.

## RESULTS

### Temporal bone - 400× magnification

At 20 and 40 days postoperatively, there were higher values in the test when compared with the control for anterior, middle, and posterior regions on both sides, with statistical significance (p<0.05). At 60 days postoperatively, there was a statistically significant difference (p<0.05) between test and control in all regions on the right side, but on the left side there was a statistically significant difference (p<0.05) between test and control, only for the middle region. The total value without stratifying into regions was always statistically different (p<0.05) between test and control on both sides (Table 1).

**Table 1.**
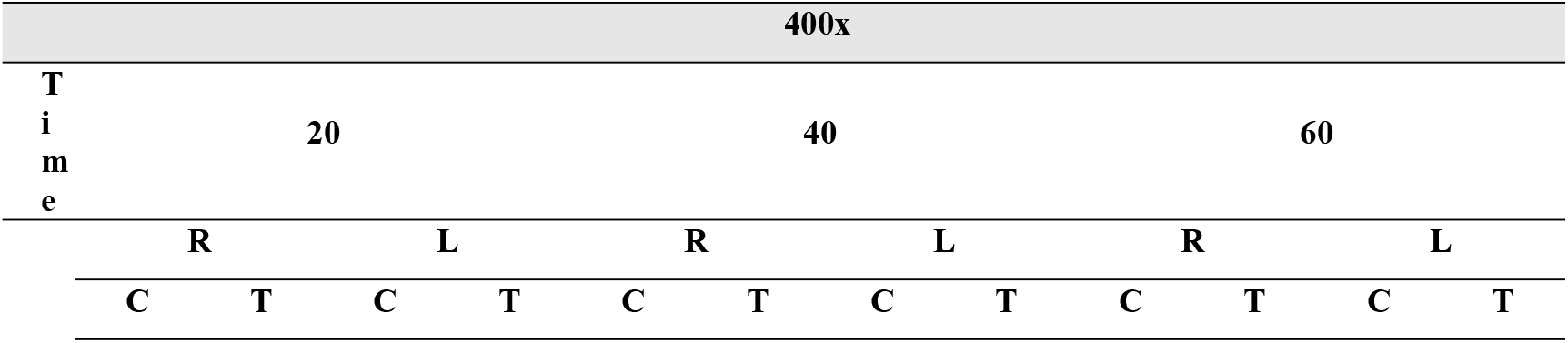
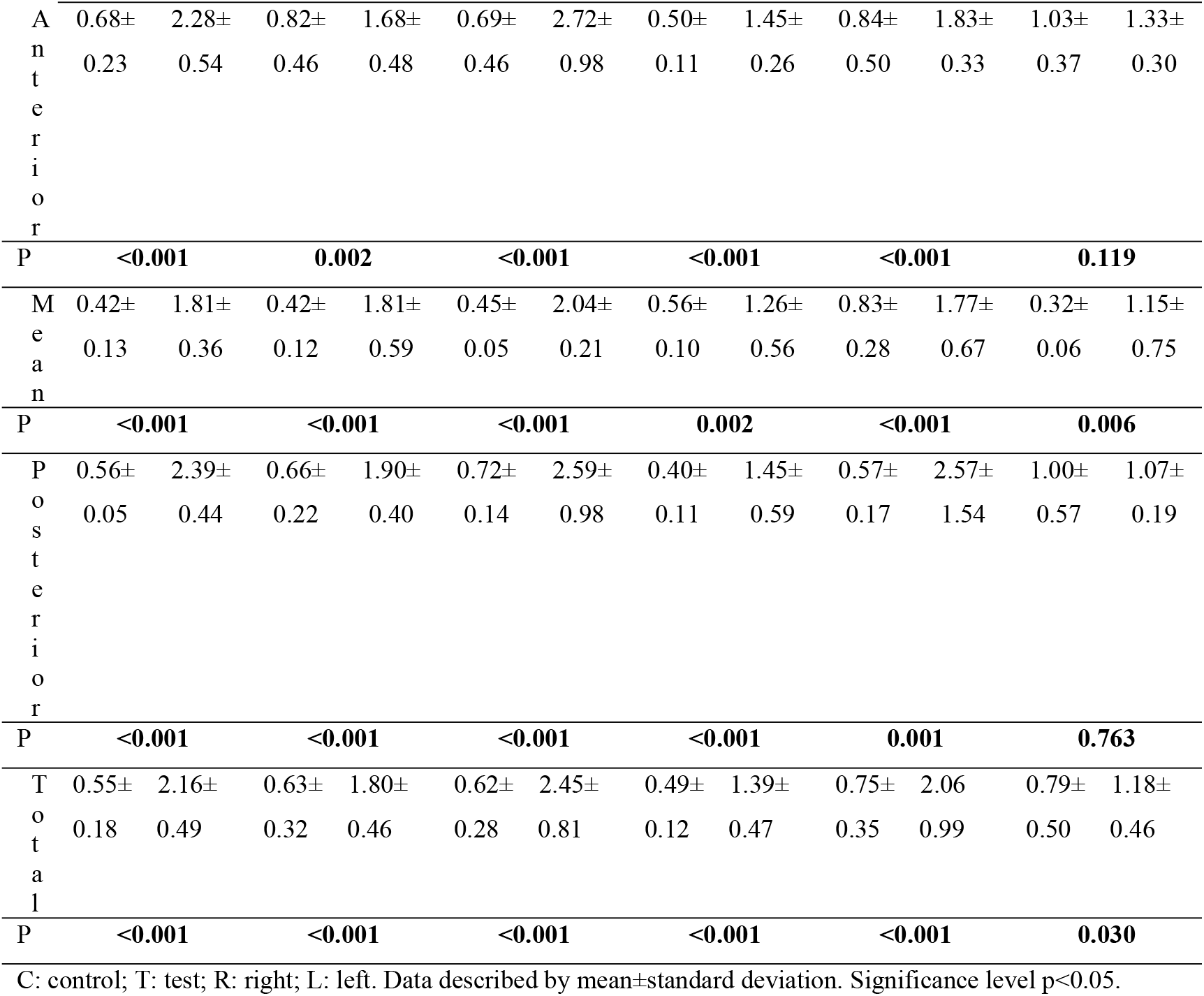
Temporal thickness in the anterior, middle, and posterior regions. Comparison between test and control groups at 20, 40, and 60 days postoperatively, for the right and left sides. Measurements in centimeters.

### Mandibular condyle - 400× magnification

At 20 and 40 days postoperatively, there were higher values in the test when compared to the control for the anterior, middle, and posterior regions on both sides, with statistical significance (p<0.05). At 60 days postoperatively, there was a statistically significant difference between the test and control in the anterior region on the right side, but the middle and posterior regions showed significance at the limit of the chosen value (5%). On the left side, there was a difference in the anterior and middle regions, but not in the posterior region. The total value without stratifying into regions was always statistically different between the test and the control on both sides (Table 2).

**Table 2.**
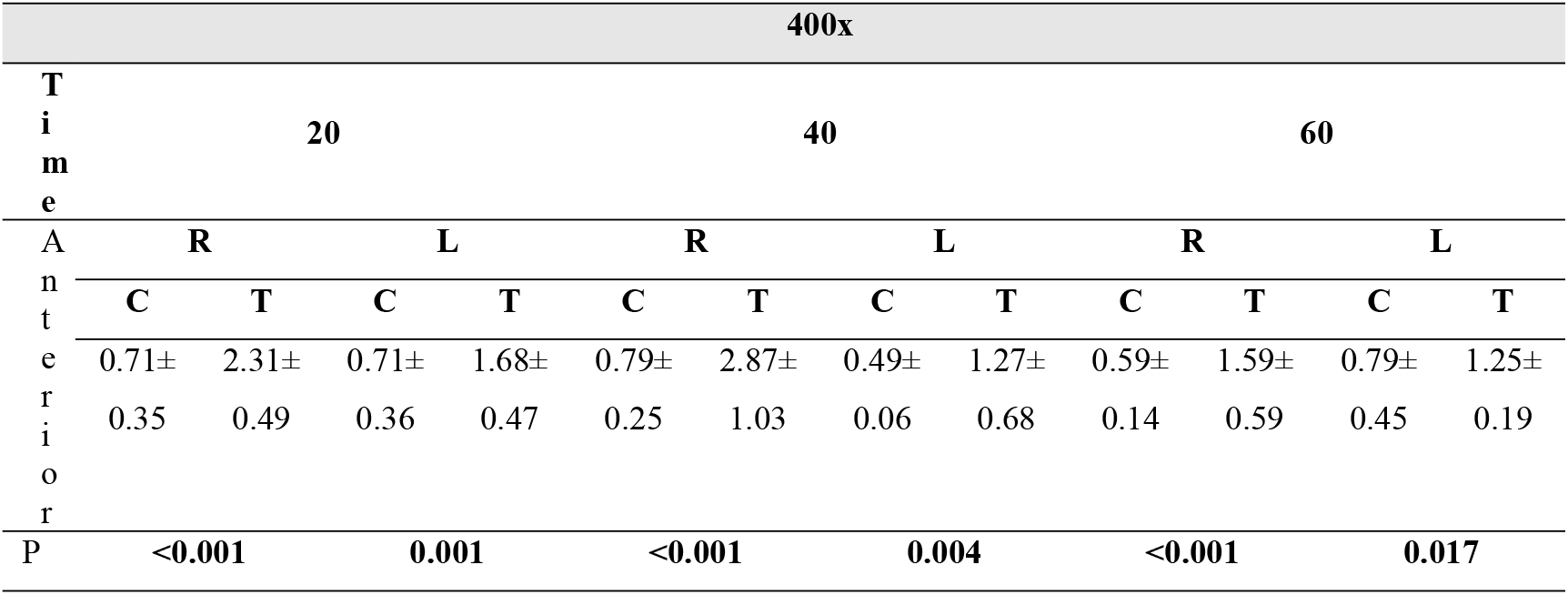
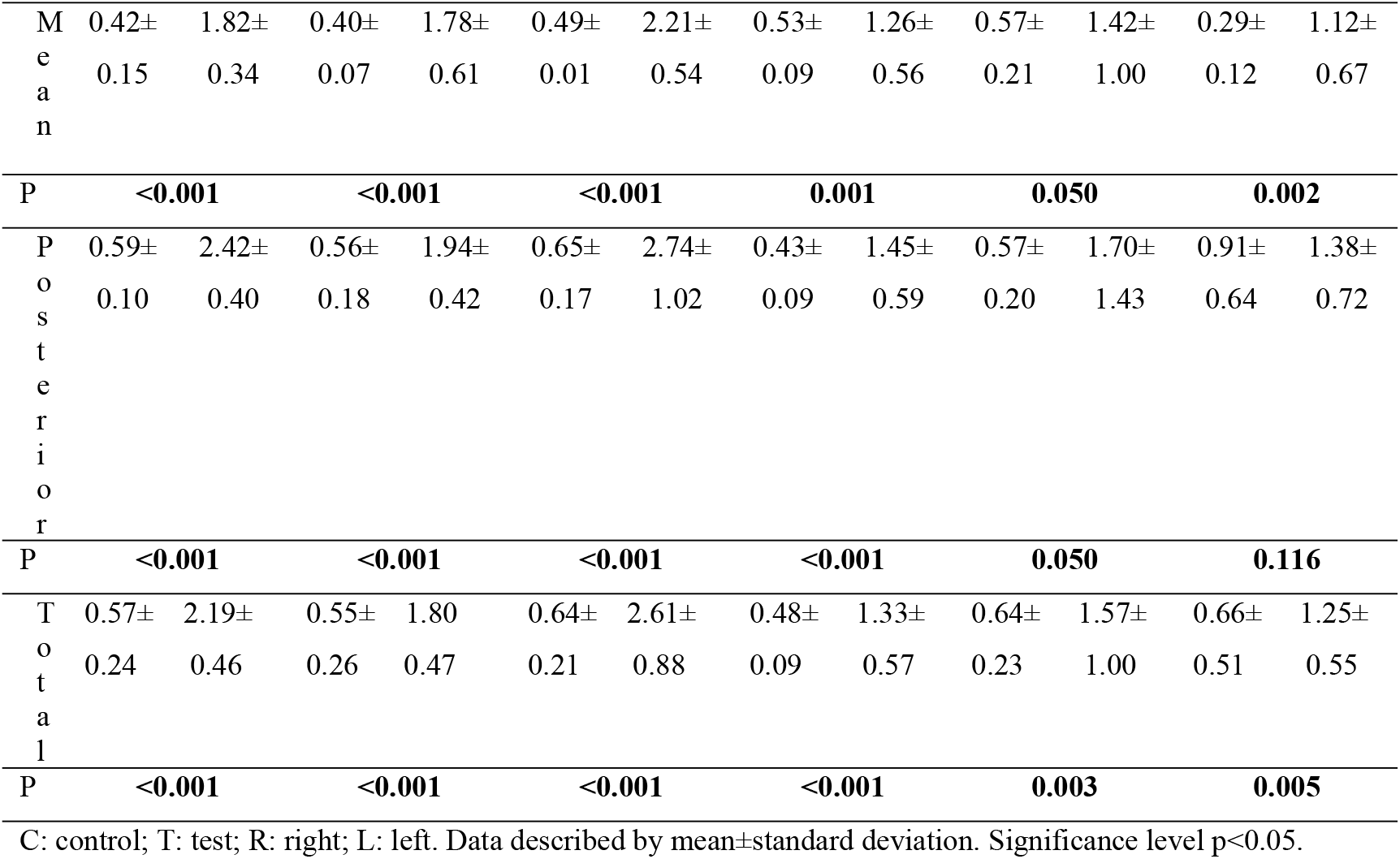
Condylar thickness in the anterior, middle, and posterior regions. Comparison between test and control groups at 20, 40, and 60 days postoperatively, for the right and left sides. Measurements in centimeters.

## DISCUSSION

The quantitative data found in this study show that there were adaptations of the temporal and condylar structures, bilaterally, considering unilateral changes in the direction of the mandibular force vector in relation to the skull base, i.e., similarly to tomographic observations in humans who underwent Puricelli Biconvex Arthroplasty. Puricelli Biconvex Arthroplasty technique proposes TMJ reconstruction by interposition of two polymethyl methacrylate (PMMA) semi-spheres. The great particularity of this technique, reproduced in this study, is the possibility of changing the direction of the force vector from the condyle, which occurs in an approach contrary to normal anatomy-physiology, since its direction turns in the sagittal inferior-superior and anterior-posterior planes [12–13].

Moreover, it is clear from the results found in this study that there is a vectorial correlation of the TMJ with craniofacial development. When the means of the control and test groups were compared under different postoperative time conditions and right and left TMJ (times 20, 40, 60; right and left sides), statistically significant differences were found in the measurements of the temporal and condylar joint surfaces. Such results confirm that the biological response to mechanical forces is the physiological property for skeletal adaptation to environmental changes [14–16]. Biomechanical factors resulting from the functional activity of the TMJ can modify the growth of its structures. Adaptation and remodeling are the main biological mechanisms responsible for maintaining the morphological and functional balance of the joint [15–19].

Ponzoni and Puricelli (2000) [18] confirmed, in another study performed in rabbits, that the change in force vector proposed by the TMJ Biconvex Arthroplasty technique provides stimulus to craniofacial growth, since induction of remodeling in the mandibular condyle was verified, as well as bone growth in the skull base. The present study performed the same force vector change as the abovementioned study, but in the current study, the histological analysis is quantitative and includes the contralateral TMJ.

Another relevant consideration of this research is the tendency to stabilize the growth of the mandibular condyle and temporal bone observed in histological analysis over the postoperative period. Although the onset of stabilization in both mandibular condyle and temporal bone was subtle, it began at 40 days postoperatively in the left side structures, intensifying at 60 days postoperatively. In this context, it should be noted that Puricelli Biconvex Arthroplasty was initially restricted to adult patients, but the indication was progressively extended to patients as young as 8 years of age, especially because it allows for a functional concept of facial growth and development [18]. From clinical analysis and CT scans, it suggests that stimulation near the tympanic fissure of the temporal bone is directly related to a sphenoid bone growth response. Another possibility to follow the postoperative evolution is by checking the occlusal function, since it allows controlling the mandibular growth and guiding the maxillomandibular relation. There are also several other studies that demonstrate that craniofacial growth can be modified, with TMJ involvement being the most important determining factor [20–22].

Some theories and hypotheses are useful to explain the nature of TMJ growth modification, with the growth relativity hypothesis and the functional matrix theory being the most advocated in the literature [23–25]. In this hypothesis, they discuss equilibrium interactions among five main factors: skeletal, viscoelastic, dental, neuromuscular, and non-muscular tissues. This set of factors contributes to TMJ adaptation by promoting increased growth of the condyle-fossa complex, redirection, and ultimately remodeling of TMJ growth [11,25].

In turn, the adaptive capacity of the TMJ articular surface corresponding to the temporal bone of rabbits is evident since its histological composition is influenced by the different characteristics of the mechanical forces that affect the TMJ during its movements and morphological changes persist in adult rabbits [15,16,18,19,26–28]. Therefore, chondrogenic responses in the temporal bone of the TMJ are more pronounced in regions where mechanical forces are higher while osteogenic responses are observed where forces are lower [29,30].

The adaptive characteristics of the rabbit TMJ show that there is a correlation between an increase in the thickness of cartilage and a decrease in the number of cartilaginous cells in response to increased forces on the surfaces of the temporal bone [12,31,32]. The remodeling process, on the other hand, occurs through osteogenesis as the bone increases in size and is characterized as a secondary process when bone reorganization occurs induced by deposition and resorption at the endosteal and periosteal surfaces [4,15,16,18,19,33].

Even though there are particularities in the comparison between human and rabbit TMJ, the morphological and histological characteristics inherent to this animal model make it the best experimental model for the proposed study. Rabbit TMJ exhibits physiological lateral and anteroposterior movements [34,35]. As a particularity of the rabbit, the articular surface of the temporal bone forms a convex eminence anteroposteriorly and concave medio-laterally. As for the articular surface of the condyle, it is known that in rabbits the anterior part is convex, both latero-laterally and anteroposteriorly [36]. The most notable morphological difference between rabbit and human TMJ(s) is the shape of the articular surface of the condyle and the retrodiscal area, since the animals do not have a post-glenoid wall [37]. Regarding histological and histochemical characteristics, similarly to the human condyle, that of the rabbit is covered by secondary cartilage and fibrous tissue [38]. In contrast, the animal cartilage is thicker in the medial region and not in the posterosuperior region. Regarding the arrangement of the cartilaginous cells, there are no significant differences between both ATM(s) [39].

Therefore, considering the surgical model used, this study confirms what has already been described in the literature, adding proof that the change in the force vector incident on the skull base promotes bilateral effects on the TMJ, demonstrating the adaptive capacity of the skeleton as a biological response to the direction of force applied.

## References

[1] Fanghänel J, Gedrange T. On the development, morphology and function of the temporomandibular joint in the light of the orofacial system. Ann Anat - Anat Anz. 2007;189:314–9. https://doi.org/10.1016/j.aanat.2007.02.024.

[2] Okeson JP. Fundamentos de oclusão e desordens temporomandibulares. 2nd ed. Artes Médicas; 1992.

[3] Monje F, Delgado E, Navarro MJ, Miralles C, Alonso Del Hoyo JR. Changes in temporomandibular joint after mandibular subcondylar osteotomy: An experimental study in rats. J Oral Maxillofac Surg. 1993;51:1221–34. https://doi.org/10.1016/S0278-2391(10)80293-9.

[4] Puricelli E, Ponzoni D, Munaretto JC, Corsetti A, Leite MGT. Histomorphometric analysis of the temporal bone after change of direction of force vector of mandible: an experimental study in rabbits. J Appl Oral Sci Science. 2012;20:526–30. https://doi.org/10.1590/S1678-77572012000500006.

[5] Tanaka Tanaka E, van Eijden T. Biomechanical Behavior of the Temporomandibular Joint Disc. Crit Rev Oral Biol Med. 2003;14(2):138–150. doi:10.1177/154411130301400207

[6] Duanmu Z, Liu L, Deng Q, Ren Y, Wang M. Development of a biomechanical model for dynamic occlusal stress analysis. Int J Oral Sci. 2021;13:29. https://doi.org/10.1038/s41368-021-00133-5.

[7] Kuroda S, Tanimoto K, Izawa T, Fujihara S, Koolstra JH, Tanaka E. Biomechanical and biochemical characteristics of the mandibular condylar cartilage. Osteoarthritis Cartilage. 2009;17:1408–15. https://doi.org/10.1016/j.joca.2009.04.025.

[8] Owtad P, Park JH, Shen G, Potres Z, Darendeliler MA. The Biology of TMJ Growth Modification. J. Dent. Res. 2013;92:315–21. https://doi.org/10.1177/0022034513476302.

[9] Nakamichi R, Asahara H. Regulation of tendon and ligament differentiation. Bone. 2021;143:115609. https://doi.org/10.1016/j.bone.2020.115609.

[10] Barnouti ZP, Owtad P, Shen G, Petocz P, Darendeliler MA. The biological mechanisms of PCNA and BMP in TMJ adaptive remodeling. Angle Orthod. 2011;81:91–9. https://doi.org/10.2319/091609-522.1.

[11] Voudouris JC, Woodside DG, Altuna G, Angelopoulos G, Bourque PJ, Lacouture CY. Condyle-fossa modifications and muscle interactions during Herbst treatment, Part 2. Results and conclusions. Am J Orthod Dentofacial Orthop. 2003;124:13–29. https://doi.org/10.1016/S0889-5406(03)00150-1.

[12] Puricelli E. Artroplastia biconvexa para tratamento da anquilose da articulação temporomandibular. Rev Fac Odontol P Alegre. 1997;38:23–7.

[13] Puricelli E. Puricelli biconvex arthroplasty as an alternative for temporomandibular joint reconstruction: description of the technique and long-term case report. Head Face Med. 2022;18–27. https://10.1186/s13005-022-00331-4.

[14] Singh M, Detamore MS. Tensile Properties of the Mandibular Condylar Cartilage. J Biomech Eng. 2008;130. https://doi.org/10.1115/1.2838062.

[15] Delpachitra SN, Dimitroulis G. Osteoarthritis of the Temporomandibular Joint: A review of Aetiology and Pathogenesis. Br J Oral Maxillofac Surg. 2021. https://doi.org/10.1016/j.bioms.2021.06.017.

[16] Sato M, Tsutsui T, Moroi A, Yoshizawa K, Aikawa Y, Sakamoto H, et al. Adaptive change in temporomandibular joint tissue and mandibular morphology following surgically induced anterior disc displacement by bFGF injection in a rabbit model. J Craniomaxillofac Surg. 2019;47:320–7. https://doi.org/10.1016/j.jcms.2018.11.034.

[17] Copray JCVM, Jansen HWB, Duterloo HS. Effects of compressive forces on proliferation and matrix synthesis in mandibular condylar cartilage of the rat in vitro. Arch. Oral Biol. 1985;30:299–304. https://doi.org/10.1016/0003-9969(85)90001-9.

[18] Ponzoni D; Puricelli E. Análise microscópica na articulação temporomandibular a partir da mudança de direção do vetor de força da mandíbula em relação à base do crânio: estudo experimental em coelhos (Oryctolagus cuniculus l). Rev Fac Odontol P Alegre. 2000;40:66–72.

[19] Fujimura K, Kobayashi S, Suzuki T, Segami N. Histologic evaluation of temporomandibular arthritis induced by mild mechanical loading in rabbits. J. Oral Pathol. Med. 2005;34:157–63. https://doi.org/10.1111/j.1600-0714.2004.00298.x.

[20] Stabrun AE, Larheim TA, Höyeraal HM, Rösler M. Reduced mandibular dimensions and asymmetry in juvenile rheumatoid arthritis. Arthritis Rheum. 1988;31:602–11. https://doi.org/10.1002/art.1780310504.

[21] Kjellberg H. Craniofacial growth in juvenile chronic arthritis. cta Odontol. Scand. 1998;56:360–5. https://doi.org/10.1080/000163598428329.

[22] Larheim TA, Haanaes HR, Ruud AF. Mandibular Growth, Temporomandibular Joint Changes and Dental Occlusion in Juvenile Rheumatoid Arthritis: *A 17-year Follow-up Study*. Scand. J. Rheumatol. 1981;10:225–33. https://doi.org/10.3109/03009748109095303.

[23] Whetten LL, Johnston LE. The control of condylar growth: An experimental evaluation of the role of the lateral pterygoid muscle. Am. J. Orthod. 1985;88:181–90. https://doi.org/10.1016/S0002-9416(85)90213-1.

[24] Perrini F, Tallents RH, Katzberg RW, Ribeiro RF, Kyrkanides S, Moss ME. Generalized joint laxity and temporomandibular disorders. J. Orofac. Pain. 1997;11:215–21.

[25] Voudouris JC, Kuftinec MM. Improved clinical use of Twin-block and Herbst as a result of radiating viscoelastic tissue forces on the condyle and fossa in treatment and long-term retention: Growth relativity. Am J Orthod Dentofacial Orthop. 2000;117:247–66.https://doi.org/10.1016/S0889-5406(00)70231-9.

[26] Hinton RJ. Changes in articular eminence morphology with dental function. Am J Phys Anthropol. 1981;54:439–55. https://doi.org/10.1002/ajpa.1330540402.

[27] Richards LC, Gurner IA. An assessment of radiographic methods for the investigation of temporomandibular joint morphology and pathology. Aust. Dent. J.p0-o 1985;30:323–32. https://doi.org/10.1111/j.1834-7819.1985.tb02524.x.

[28] Tuominen M, Kantomaa T, Pirttiniemi P. Effect of food consistency on the shape of the articular eminence and the mandible: An experimental study on the rabbit. Acta Odontol. Scand. 1993;51:65–72. https://doi.org/10.3109/00016359309041150.

[29] Silbermann M, Lewinson D, Gonen H, Lizarbe MA, von der Mark K. In vitro transformation of chondroprogenitor cells into osteoblasts and the formation of new membrane bone. Anat. Rec. 1983;206:373–83. https://doi.org/10.1002/ar.1092060404.

[30] Tallents RH, Macher DJ, Rivoli P, Puzas JE, Scapino RP, Katzberg RW. Animal model for disk displacement. J Craniomandib Disord. 1990;4:233–40.

[31] McCormick SU, McCarthy JG, Grayson BH, Staffenberg D, McCormick SA. Effect of Mandibular Distraction on the Temporomandibular Joint. J. Craniofac. Surg. 1995;6:358–63. https://doi.org/10.1097/00001665-199509000-00005.

[32] Puricelli E. Cirugía bucomaxilofacial en el paciente pediatrico. In: Villa CN, Marín FG, Caicoya SO Tratado de Cirugía Oral y Maxilofacial. 2009:1879–901.

[33] Enlow D. Facial Growth and Development. Int J Orofacial Myology. 1979;5:7–10. https://doi.org/10.52010/ijom.1979.5.4.3.

[34] Fox SS. Lateral jaw movements in mammalian dentitions. J Prosthet Dent. 1965;15:810–25.

[35] Weijs WA, Brugman P, Grimbergen CA. Jaw movements and muscle activity during mastication in growing rabbits. Anat. Rec. 1989;224:407–16. https://doi.org/10.1002/ar.1092240309.

[36] Mills DK, Daniel JC, Scapino R. Histological features and in-vitro proteoglycan synthesis in the rabbit craniomandibular joint disc. Arch. Oral Biol. 1988;33:195–202. https://doi.org/10.1016/0003-9969(88)90045-3.

[37] Kyllar M, Putnová B, Jekl V, Stehlík L, Buchtová M, Štembírek J. Diagnostic imaging modalities and surgical anatomy of the temporomandibular joint in rabbits. Lab. Anim. 2018;52:38–50. https://doi.org/10.1177/0023677217702178.

[38] Mizoguchi I, Takahashi I, Nakamura M, Sasano Y, Sato S, Kagayama M, et al. An immunohistochemical study of regional differences in the distribution of type I and type II collagens in rat mandibular condylar cartilage. Arch. Oral Biol. 1996;41:863–9. https://doi.org/10.1016/S0003-9969(96)00021-0.

[39] Al-Moraissi EA, Wolford LM. Does Temporomandibular Joint Pathology With or Without Surgical Management Affect the Stability of Counterclockwise Rotation of the Maxillomandibular Complex in Orthognathic Surgery? A Systematic Review and Meta-Analysis. J. Oral Maxillofac. Surg. 2017;75:805–21. https://doi.org/10.1016/j.joms.2016.10.034.

